# Subjective Audibility Modulates the Susceptibility to Sound-Induced Flash Illusion

**DOI:** 10.1101/2022.06.22.497147

**Authors:** Yuki Ito, Hanaka Matsumoto, Kohta I. Kobayasi

**Affiliations:** Organization for Research initiatives and Development, Doshisha University, 1-3 Tataramiyakodani, Kyotanabe, 610-0394, Japan; Graduate School of Life and Medical Sciences, Doshisha University, 1-3 Tataramiyakodani, Kyotanabe, 610-0394, Japan

**Keywords:** audio-visual integration, multisensory, sound-induced flash illusion, subjective audibility

## Abstract

When a brief flash is presented along with two brief sounds, the single flash is often perceived as two flashes. This phenomenon is called a sound-induced flash illusion, in which the auditory sense, with its relatively higher reliability in providing temporal information, modifies the visual perception. Decline of audibility due to hearing impairment is known to make subjects less susceptible to the flash illusion. However, the effect of decline of audibility on susceptibility to the illusion has not been directly investigated in subjects with normal hearing. The present study investigates the relationship between audibility and susceptibility to the illusion by varying the sound pressure level of the stimulus. In the task for reporting the number of auditory stimuli, the lowering of the sound pressure level caused the rate of perceiving two sounds to decrease on account of forward masking. The occurrence of the illusory flash was reduced as the intensity of the second auditory stimuli decreased, and was significantly correlated with the rate of perceiving the two auditory stimuli. These results suggest that the susceptibility to sound-induced flash illusion depends on the subjective audibility of each sound.

## 1. Introduction

We utilize information from multiple modalities when perceiving the world. In most cases, integrating the auditory and visual senses reduces the ambiguity of unimodal information and aids in reconstructing a reliable perceptual world. The sensory integration usually improves behavioral tasks, such as stimulus detection (Bolognini *et al*., 2005; Frassinetti *et al*., 2002) and speech perception (Ross *et al*., 2007; Sumby and Pollack, 1954). However, in some cases, when we receive conflicting information from different modalities the perception of one sensory modality is significantly affected by the other, creating illusions such as the Ventriloquism effect (Jack and Thurlow, 1973), McGurk effect (McGurk and MacDonald, 1976) and the Stream-bounce effect (Sekuler *et al*., 1997).

Recently, sound-induced flash illusion, whereby, a single flash is perceived as two, when two brief sounds are simultaneously presented, has attracted the attention of many scientists as this illusion is induced by a simple set of stimulus and is highly replicable (Shams *et al*., 2002; Shams *et al*., 2000). It is proposed that sound-induced flash illusion follows the optimal rule of multisensory integration, whereby, the weighting of a sensory modality depends on its reliability and precision, and the more reliable modality dominates the final perception (Shams *et al*., 2005). Audition is relatively strongly weighted than vision during the perception of fast-changing stimulus, and has a larger impact on the final perception as the auditory modality has a better temporal resolution than the visual modality (Welch and Warren, 1980). In addition, a causal inference model, which takes into account the prior knowledge on the world including the usefulness of each modality, predicated that a person’s sensitivity to perceive auditory and visual stimuli mediates their susceptibility to the sound-induced flash illusion. Several studies on the auditory-impaired or age-related sensory declined subjects partially support the idea. For example, the susceptibility to sound-induced flash illusion among hearing aid users is stronger than in hearing-impaired non-users (Gieseler *et al*., 2018), and the susceptibility to the illusion is positively correlated with the self-reported hearing ability in aged (> 50 years old) subjects (Hirst *et al*., 2019). These researches demonstrate that degradation in sound audibility due to sensory decline makes the subject less susceptible to the illusion. However, the effect of degradation of audibility due to sound quality on susceptibility to the illusion has not been systematically investigated in normal-hearing subjects.

One of the most commonly observed and well-studied phenomena interfering with the audibility of sound is temporal masking (Moore and Glasberg, 1983). Perception of a sound is interfered when another louder sound is presented right before (< 200ms) the sound, and this is called forward masking (Moore, 2012). The audibility of the sound decreases as the sound pressure level of the signal is lowered, and it eventually becomes totally undetectable. In this study, we tried to systematically modulate the audibility of sound stimulus by utilizing forward masking. Specifically, a visual stimulus, a brief flash or a pair of flashes, was presented with a pair of tone bursts, where the intensity of the second sound varied while the intensity of the first sound was kept constant. The effect of masking on the illusion was examined in experiment I, and the relationship between the masking effect and the susceptibility to the sound-induced flash illusion was evaluated in experiment II.

## 2. Materials and Methods

### 2.1 Participants

Twenty adults (11 women and 9 men, 21-29 years old) participated in this experiment, and each participant took part in only one of the two experiments. Ten individuals (4 women and 6 men) participated in experiment I, and the remaining ten (7 women and 3 men) participated in experiment II. All participants have normal hearing and normal, or corrected normal, vision. Informed consent was obtained from all participants before experiments. All experiments were conducted in accordance with the guidelines for human experiments approved by the Ethics Committee of Doshisha University.

### 2.2 Stimuli

The visual stimulus was a uniform white disk (55 mm diameter) displayed on a black background using a liquid crystal display (532 mm width x 295 mm height, Foris FG2421, Eizo). A fixation point (6 mm white cross) was displayed at the center of the screen throughout the entire session, and the white disk was presented at 21 mm below the fixation point for 8 ms. The visual stimulus was presented in two ways: flashing once (single flash) or twice (double flashes). The inter-stimulus interval (ISI) between the offset of the first stimulus and the onset of the second stimulus was set at 58 ms when double flashes were presented (Fig. 1). The auditory stimulus was a brief tone burst with a frequency of 3 kHz, and the duration of the auditory stimulus was 20 ms (the rise and fall time were 5 ms). The auditory stimulus was always presented twice with an ISI of 48 ms via headphones (SR-507, STAX). In experiment I, two stimulus settings for sound pressure level were used: a same intensity condition and a different intensity condition. The sound intensity of the first and second sounds was identical in the same intensity condition. The sound pressure level of the stimulus was 70, 80, and 90 dB SPL. In the different intensity condition, the sound pressure level of the first stimulus was set at 80 dB SPL, and the intensities of the second sounds varied between 70, 73, 75, 78, 82, 85, 87, and 90 dB SPL.

**Fig. 1.**
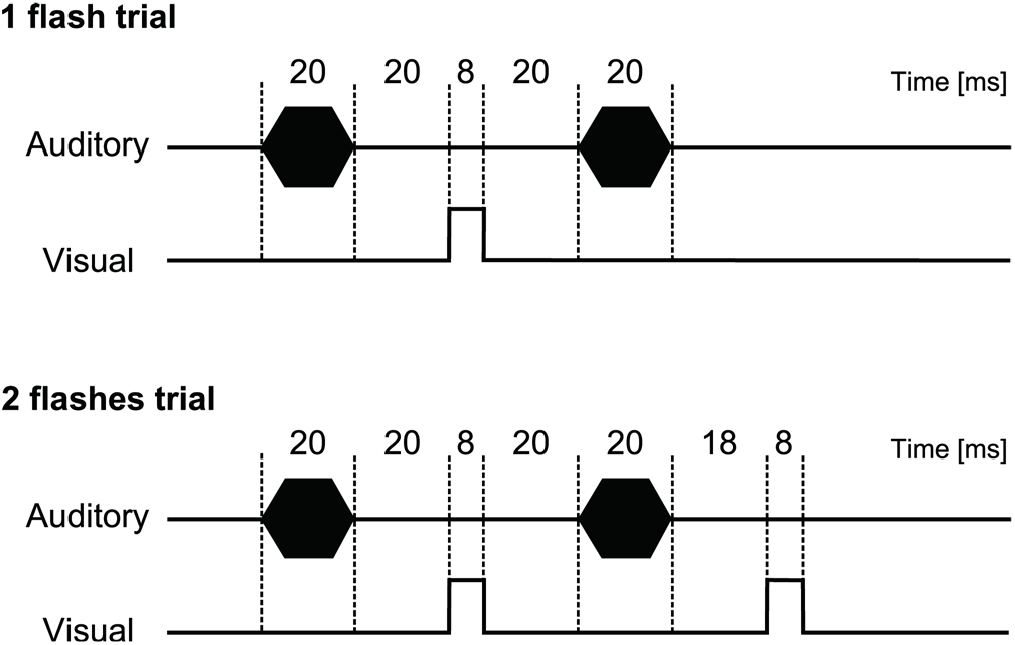
Temporal configuration of stimuli. The stimulus consisted of two-tone bursts, and either one flash or two flashes. The durations of the auditory and visual stimuli were 20 ms and 8 ms, respectively. The first tone burst was presented 40 ms before the onset of a flash. The two sounds were separated by 48 ms and flashes by 56 ms.

Experiment II consisted of a visual task and an auditory task. In the visual task trial, the participants were asked to report the number of visual stimuli they observed, and in the auditory task trial, they were asked to report the number of auditory stimuli sensed. Auditory stimulus having 75 dB SPL was always presented as the first tone burst. The sound pressure level of the second sounds were at 55, 59, 63, 67, 71, 75, 79, 83, 87, 91, and 95 dB SPL in the visual task, and 55, 63, 75, 87, and 95 dB SPL in the auditory task. The first tone burst of a tone pair preceded the single flash and the first flash of a flash pair by 40 ms. The refresh rate of the monitor was set at 120 Hz, and the duration of visual stimulus was confirmed by a high-speed camera (EX-F1, Casio) with a frame rate of 1000 per second. The sound pressure level was measured and adjusted with a microphone (ER-7C Series b, Etymotic research).

### 2.3 Task and Procedure

The experimental procedure was the same as our previous research (Ito *et al*., 2020). The participants sat in front of a monitor placed 60 cm from their faces, and the height of the monitor was adjusted so that the position of their eye was levelled with the fixation point. In experiment I, participants were instructed to state the number of flashes they perceived by pressing a button on a keyboard. After they had answered, the next trial began automatically after a 1 s interval. Each stimulus pair (2 types of flashes × 11 sounds) was presented 11 times in random order, resulting in 242 trials for each participant. In experiment II, the participants were asked to state the number of flashes (Visual trial) or sounds (Auditory trial) they perceived. They were blind to which task they were in during stimulus presentation, and were informed of the task immediately after the stimulus was presented. We instructed the participants to focus on the visual stimuli. The participants completed a total of 309 trials. There were 12 trials for each stimulus pair (2 types of flashes × 11 sounds) in the visual task, and 9 trials for each sound (5 sounds) in the auditory task. The order of the stimuli and tasks was randomized. All experiments were controlled with the Presentation software (Neurobehavioral Systems) and were conducted in a sound-proof room (170-cm width × 150-cm length × 230-cm height). The noise level at the sound-proof room was below 38.3 dB SPL.

## 3. Results

### 3.1 Experiment I

When the sound pressure level of the first and second sound were the same (same intensity condition), the average percentage of double-flashes perception in the trial of 90 dB SPL was marginally higher than that in the trial of 70 dB SPL (42.7 ± 9.1 % vs. 49.1 ± 9.0 % in 1 flash trial; 70.9 ± 8.3 % vs. 80.9 ± 4.6 % in 2 flashes trial, mean ± S.E.M.; Fig.2A). A two-way ANOVA with flash and intensity as within-subject factors was performed. The main effect of flash type was significant *(F*_(1,9)_ = 15.87, *p* < 0.01, *η*_*p*_^2^ = 0.64), while a significant main effect of intensity (*F*_(2,18)_ = 1.89, *p* = 0.18, *η*_*p*_^2^ = 0.17) and flash type × intensity interaction (*F*_(2,18)_ = 0.99, *p* = 0.39, *η*_*p*_^2^ = 0.10) was not observed.

In the different intensity condition in which the sound pressure level of the first auditory stimulus was different from that of the second auditory stimulus, the rate of perception of two flashes increased as the intensity of the second tone burst increased in both single and two flashes condition (Fig. 2B). A two-way ANOVA with flash and intensity revealed a significant main effect of flash type (*F*_(1,9)_ = 15.69, *p* < 0.01, *η*_*p*_^2^ = 0.64) and intensity (*F*_(8,72)_ = 9.01, *p* < 0.001, *η*_*p*_^2^ = 0.50). There was no significant interaction between flash and intensity (*F*_(8,72)_ = 0.59, *p* = 0.78, *η*_*p*_^2^ = 0.06).

**Fig. 2.**
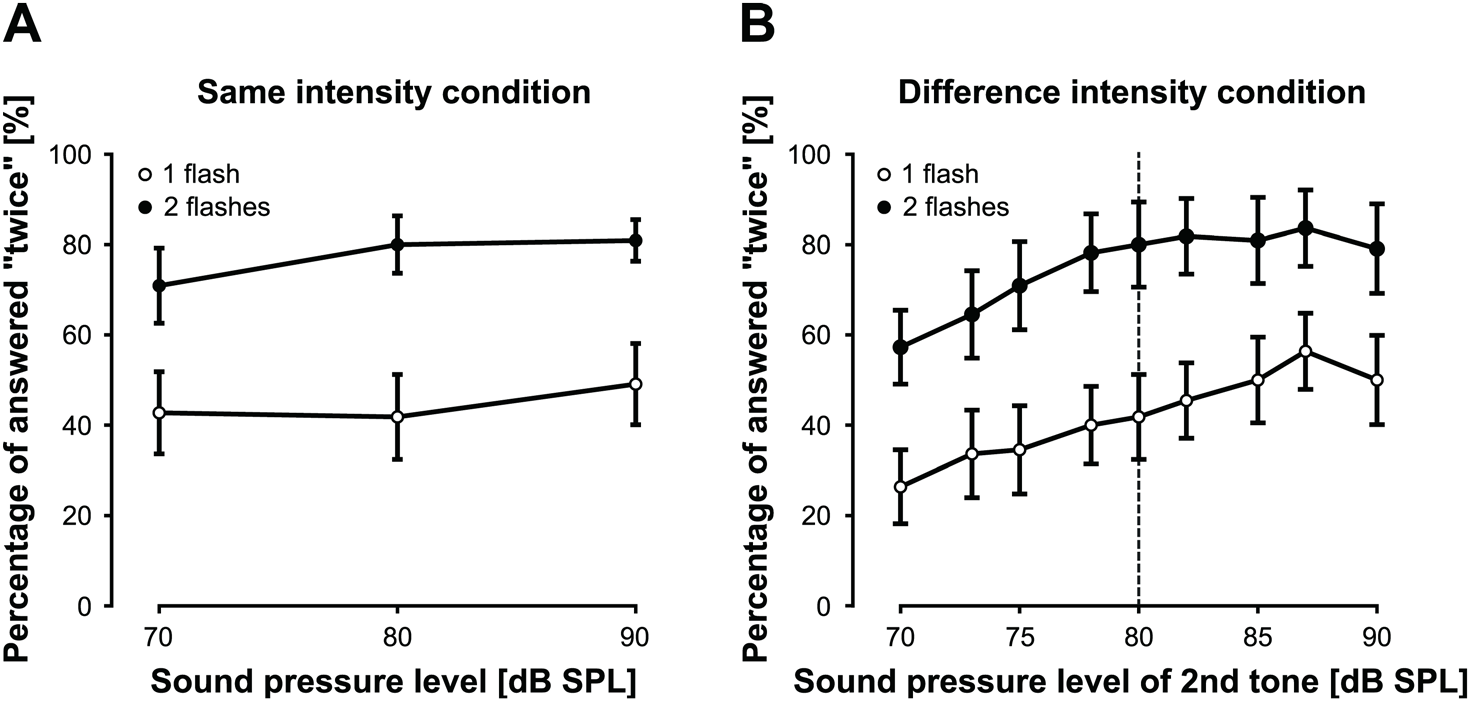
Mean proportion of perception of the two flashes in the same (A) and different intensity (B) condition. (A) In the same intensity condition the sound pressure level of the first and second auditory stimulus was the same. The open circle indicates the results of the one-flash trial, and the closed circle indicates that of the two-flashes trial. (B) In the different intensity condition the intensity of the first and second auditory stimulus were different. The sound pressure level of the first tone burst was always 80 dB SPL, which was denoted by a vertical dashed line. Error bars indicate the standard error of the mean.

### 3.2 Experiment II

In the visual task, the rate of perception of the two flashes increased as the sound pressure level of the second auditory stimulus increased both in single and double flash trials (Fig. 3A). A two-way ANOVA with flash and intensity showed a significant effect of flash (*F*_(1,9)_ = 14.79, *p* < 0.01, *η*_*p*_^2^ = 0.62) and intensity (*F*_(10,90)_ = 12.69, *p* < 0.001, *η*_*p*_^2^ = 0.59); however, no significant effect of flash × intensity interaction (*F*_(10,90)_ = 1.58, *p* = 0.13, *η*_*p*_^2^ = 0.15) was observed. In the auditory task, a one-way ANOVA on the number of perceived flashes revealed a clear significant effect of the sound pressure level (*F*_(4,36)_ = 13.72, *p* < 0.001, *η*_*p*_^2^ = 0.60). The rate of perception of the two sounds increased as the intensity of the second tone burst increased (Fig. 3B). In addition, we investigated the relationship between the occurrence of the illusion and the perception of paired auditory stimulus with repeated measures correlation (Bakdash and Marusich, 2017). In a single flash trial, repeated measures correlation analysis revealed a significant positive correlation between the percentage of perception of the two sounds and the occurrence of illusory flash (*r*_*rm*_(39) = 0.79, 95% CI [0.63, 0.89], *p* < 0.001; Fig. 4A). Similarly, significant repeated measures correlation between the rate of perception of the two sounds and the rate of perception of two flashes was observed (*r*_*rm*_(39) = 0.80, 95% CI [0.66, 0.89], *p* < 0.001; Fig. 4B).

**Fig. 3.**
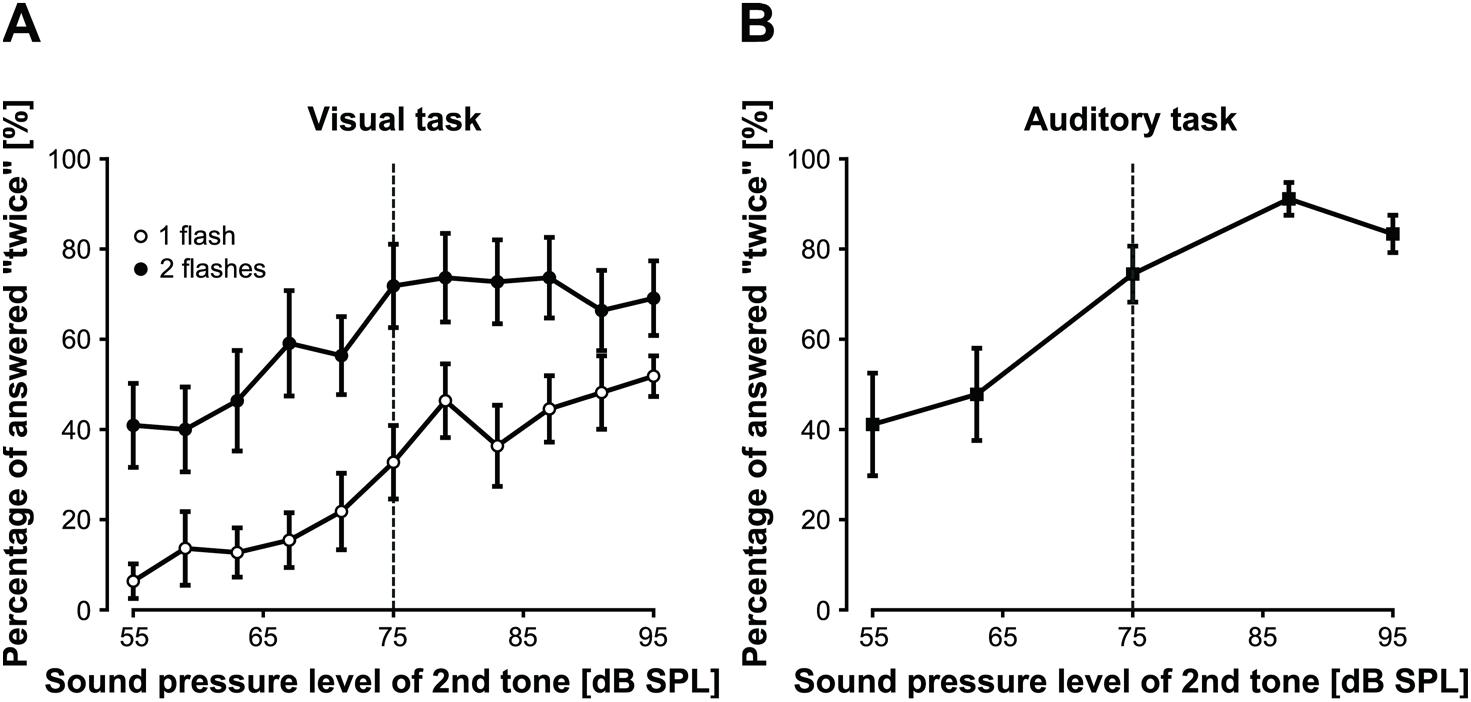
Mean proportion of perception of the two flashes in visual task (A) and two tone perception in auditory task (B). The two auditory stimuli with different sound pressure level were presented. The intensity of the first tone burst was always 75 dB SPL, which was denoted by a vertical dashed line. (A) As the sound pressure level of the second sound became lower than that of the first sound, the fusion illusion occurred more frequently; however, the fission illusion occurred less. (B) The forward masking decreased the audibility of the second sounds. Error bars indicate the standard error of the mean.

**Fig. 4.**
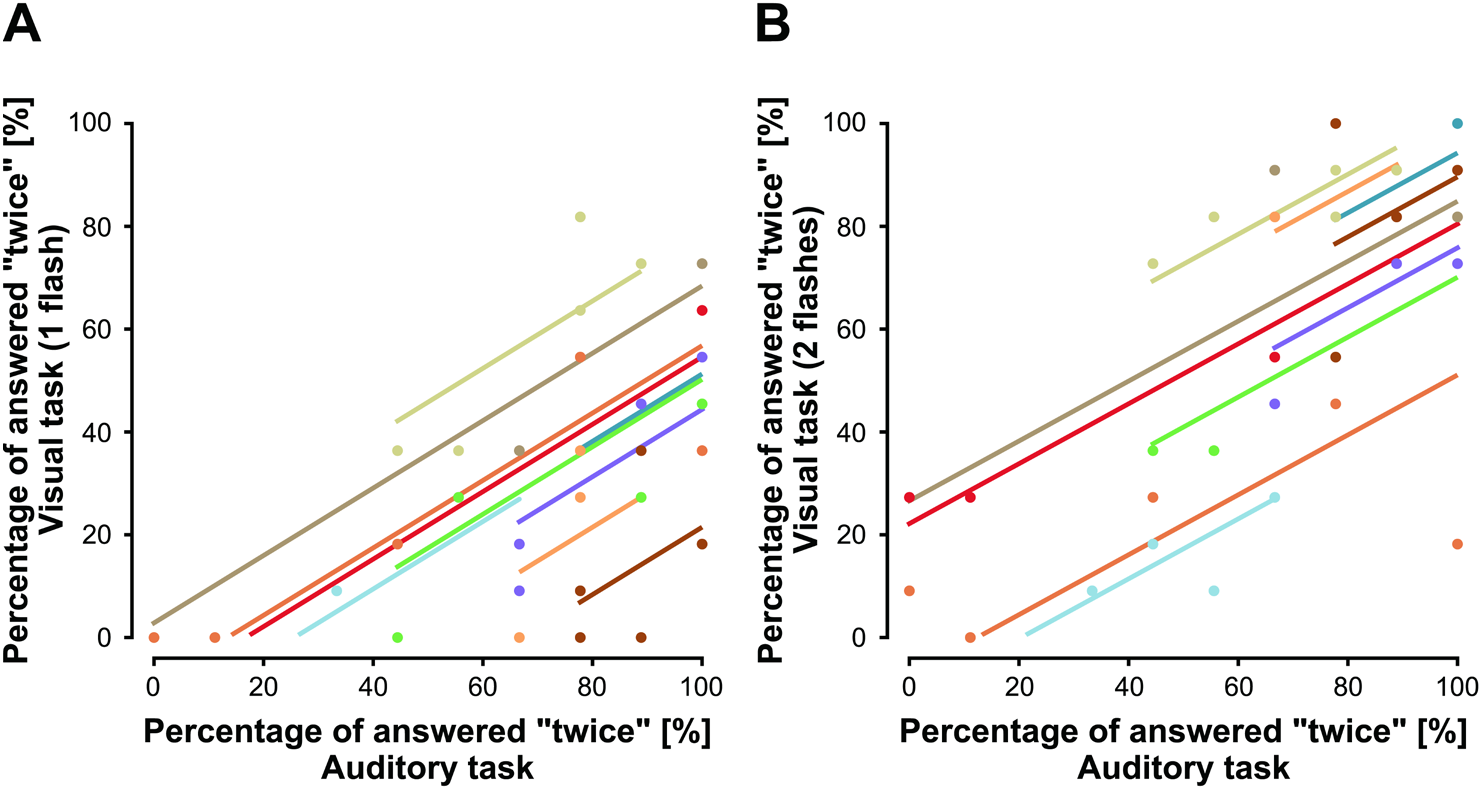
Correlations between audibility and the susceptibility to perceive two flashes. Each dot shows one of five sound intensity configurations (same as depicted in Fig. 3B) from each individual. The colored lines demonstrate the regression for the within-subject correlation. The abscissa represents the rate of the auditory stimulus perceived twice in auditory task, and the ordinate represents the rate of the visual stimulus perceived twice in visual task. For both the single flash (A) and double flash (B) trial, the two-flashes perception increased as the two-sounds perception increased. Repeated-measures correlation analysis showed a significant positive correlation between the two (*r*_*rm*_(39) = 0.79, *p* < 0.001 in one flash trial; *r*_*rm*_(39) = 0.80, *p* < 0.001 in two flashes trial).

## 4. Discussion

In this study, we examined whether the subjective perception of sounds affected the occurrence of sound-induced flash illusion by varying the sound pressure level of the auditory stimuli. Experiment I revealed that it became difficult to induce the double flash illusion when the sound pressure level of the second auditory stimuli was lower than that of the first sound (Fig. 2B). Experiment II revealed that the occurrence of the illusion was correlated with the audibility of the two sounds (Fig. 4).

In same intensity condition in experiment I, when tone burst of high intensity was presented, the percentage of perceiving two flashes was slightly higher than when low-intensity stimuli were presented (Fig. 2A). Andersen *et al*. (2004) showed that the illusory flash did not occur when the intensity of auditory stimuli was near the threshold level. Our data is partially consistent with their result in terms of intensity dependence; however, the effect was not significant. The sound pressure level of all sounds used in this research were sufficiently loud (i.e. > 70 dB SPL) and the intensity dependence could plateau at this level. In different intensity condition in experiment I, when sound pressure level of the second tone burst was not the same as the first tone burst (Fig. 2B), our data showed that the occurrence of illusory flash decreased as the second tone intensity lowered. The result of ANOVA shows that the rate of two flashes perception significantly changes depending on the stimulus intensity in different intensity condition (Fig. 2B), while no significant effect of amplitude is observed in same intensity conditions (Fig. 2A). Therefore, the variation of illusory flash perception in different intensities is not due to absolute sound pressure levels but rather due to intensity deference.

When two sounds are consecutively presented with a short inter-stimulus interval (ISI), we often miss the presence of the second stimulus (Moore and Glasberg, 1983); and the effect is called a temporal masking. The amount of masking increases as the intensity of the first stimulus (masker) increases (Moore and Glasberg, 1983). Forward masking occurs when the ISI between auditory stimuli is within 200 ms, while backward masking is little seen when the ISI exceeds 25 ms (Durrant *et al*., 1985). Since ISI between auditory stimuli used in this study were 48 ms, the audibility of the second were affected mostly by forward masking. However, it was unclear in experiment I whether the subjective audibility was altered by forward masking. Accordingly, we conducted experiment II to confirm that the second sound was harder to perceive as the intensity of the sound decreases in our stimulus settings. Specifically, the auditory task, in which participants tried to identify the number of auditory stimuli, was conducted together with the visual task, which was identical to the different intensity trials of experiment I. In addition, we tested widened intensity difference between the first and second sound as we expected the effect of masking to be more pronounced.

Consistent with the result of experiment I, the susceptibility of flash illusion decreased in the visual task (Fig. 3A) as the sound pressure level of the second tone burst weakened. As expected, the rate at which participants perceived the two sounds decreased as the sound pressure level of the second sound decreased (Fig. 3B). In addition, repeated measures correlation analysis shows significant in-subject correlations between the audibility of the two sounds and the perception of the two flashes (Fig. 4). These results indicate that the occurrence of the double flash illusion decreased when the audibility of the sound was reduced due to forward masking; therefore, the two sounds need to be clearly heard for perceiving the illusory second flash.

While we have so far focused on the fission illusion, in which one flash accompanied with two sounds induces the perception of two flashes, previous studies have also observed fusion illusion, whereby the double flashes were perceived as single when accompanied with one sound (Andersen *et al*., 2004; Shams *et al*., 2005). Notably, in the double flashes trial, in which the visual stimulus was physically presented twice, the rate of perception of the two flashes decreased as the intensity of the second auditory stimulus decreased in both experiment I (Fig. 2B) and II (Fig. 3A). The decrease in the perception of the two flashes can be attributed to the fusion illusion, as forward masking caused participants to perceive the two sounds as one. Our data showed that the occurrence of both the fission and fusion illusion is modulated in a consistent manner by regulating the sound pressure level of the second auditory stimulus.

Many studies have observed that sound-induced flash illusion is robust in the feature of visual stimulus (e.g. Shape: Takeshima, 2020; Gabor patch: Takeshima and Gyoba, 2015, Block pattern: Takeshima and Gyoba, 2013; Gaussian probs: Apthorp *et al*., 2013; Faces: Setti and Chan, 2011). However, despite sound being the cause of the phenomenon, few studies have investigated the effect of features of auditory stimulus on this illusion. Roseboom *et al*. (2013) revealed that the sound-induced flash illusion did not occur when the two auditory stimuli had distinct frequency difference. In addition, our previous research demonstrated that the susceptibility of the fission illusion decreased as a function of the frequency difference between two auditory stimuli (Ito *et al*., 2020). These studies suggest that the similarity between the two auditory stimuli promote the occurrence of illusory flash. Our results, however, partially contradict this concept. When the second sound became louder than the first, the fission illusion continued to occur even though the intensity differences widened (Fig. 3A). This result suggests that dissimilarity in the amplitude does not always inhibit the occurrence of illusory flash and shows clear contrast with the sensitivity to frequency difference, in which a 1/3 octave frequency difference significantly inhibits the illusion (Ito *et al*., 2020). Additionally, it provides evidence for the hypothesis that frequency correspondence of auditory stimuli is an important factor in the occurrence of sound-induced flash illusion.

In conclusion, the present study investigated how the subjective audibility of the sound modulate the susceptibility to the sound-induced flash illusion (both fission and fusion). The second sound intensity was modified to regulate the forward masking and thereby control audibility of the second sound. The occurrence of the fission illusion increased and the fusion illusion decreased, when a louder second sound was presented. These results were consistent with between-subject studies of age-related hearing decline (Hirst *et al*., 2019) and hearing aid users (Gieseler *et al*., 2018), which showed that the susceptibility of fission was reduced as the audibility decreased. Our data expand the knowledge on normal hearing subjects and suggests that susceptibility to the illusion is modulated by the audibility of stimulus within the subject.

## Acknowledgments

This research was supported by the JSPS KAKENHI (19J22981 to YI, 21H03469 to KIK).

## Notes

### Competing Interest Statement

The authors have declared no competing interest.

